# Hailey-Hailey disease models identify synergistic therapeutic effects of MEK and ROCK inhibition

**DOI:** 10.64898/2026.05.20.726679

**Authors:** Jessica L. Ayers, Arti Parihar, Afua Tiwaa, Akshata Aravind, Michael C. Martin, Kuga Pence, Crystal J. Tam, Nizhoni Sutter, Kristen Skruber, Mrinal K. Sarkar, Johann E. Gudjonsson, Cory L. Simpson

**Affiliations:** Department of Dermatology, University of Washington, Seattle, WA, USA; Department of Laboratory Medicine & Pathology, University of Washington, Seattle, WA, USA; Department of Biology, Gonzaga University, Spokane, WA, USA; Department of Dermatology, University of Michigan, Ann Arbor, MI, USA; Institute for Stem Cell & Regenerative Medicine, University of Washington, Seattle, WA, USA

## Abstract

Hailey-Hailey disease (HHD) is a genetic skin blistering disorder lacking approved treatments despite linkage to *ATP2C1* variants 25 years ago. Since knockout mice did not replicate HHD, we ablated *ATP2C1* in human keratinocytes or chemically inhibited its encoded Golgi calcium pump SPCA1. In organotypic epidermis, SPCA1 deficiency or inhibition reproduced HHD pathology, disrupting desmosomal cadherins and severing cell-cell junctions, termed acantholysis. RNA sequencing of heterozygous cells identified dysregulation of actin and Rho GTPases along with EGF receptor signaling as potential pathogenic drivers. Accordingly, SPCA1-depleted organotypic epidermis and HHD biopsies exhibited cortical actin disorganization and hyper-phosphorylation of the Rho kinase (ROCK) target, myosin light chain. Rho activation was sufficient to induce acantholysis, while ROCK inhibition partially restored heterozygous keratinocyte cohesion. A fluorescent biosensor demonstrated ERK hyper-activation in heterozygous cells along with desmosomal cadherin mis-localization. Importantly, treating SPCA1-deficient keratinocyte sheets with MEK and ROCK inhibitors together fully restored their integrity. Our results show HHD blistering is driven by desmosome and cortical actin dysfunction that was mitigated by targeting MEK and ROCK with repurposed drugs, offering a viable treatment strategy. Moreover, our model provides a blueprint for replicating genetic epidermal disorders to delineate pathogenic mechanisms and vet therapeutics for other orphan skin diseases.

**GRAPHICAL ABSTRACT:** 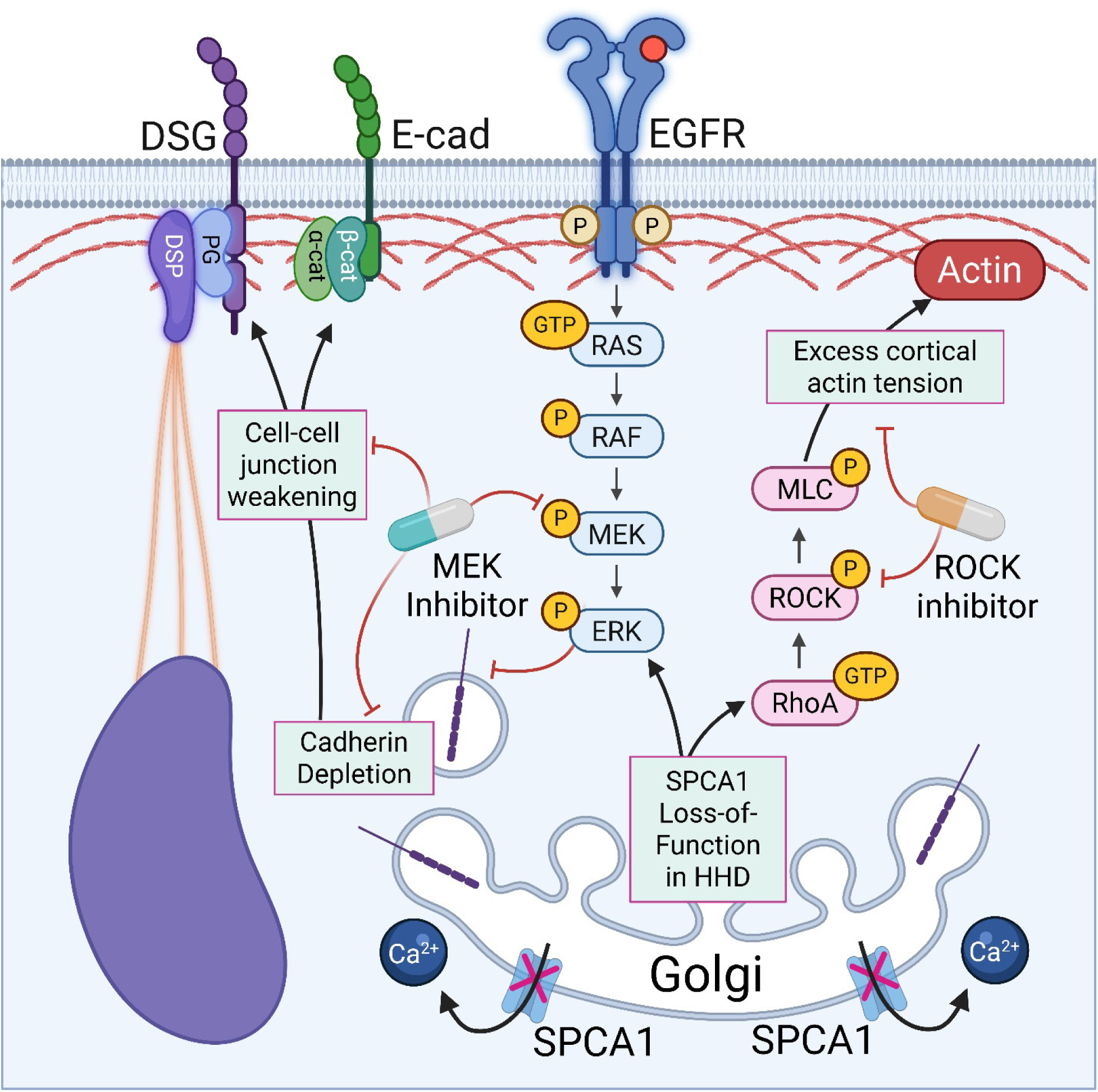

## INTRODUCTION

Hailey-Hailey disease (HHD), originally called familial benign chronic pemphigus [1] and recently termed *ATP2C1* non-syndromic epidermal differentiation disorder (*ATP2C1*-nEDD) [2], is an orphan skin blistering disease [3]. It is characterized by impaired keratinocyte adhesion [4], which causes epidermal blisters and painful skin fissures most often in body folds like the axillae and groin [5, 6]. Patients who suffer from HHD endure a lifetime of repeated flares, social isolation, disability, and reduced quality of life [7]. Breakdown of the epidermal barrier in HHD permits recurrent skin super-infections [8, 9], in severe cases leading to hospitalization or even death [10].

HHD is caused by dominant variants in *ATP2C1* [11, 12], leading to haplo-insufficiency of its encoded P-Type ATPase, SPCA1 [13]. This ion pump localizes to the Golgi and is responsible for moving cytoplasmic calcium into its lumen [14], the site for critical post-translational modifications of extracellular protein domains. To mediate adhesion, cadherin ectodomains must undergo pro-domain cleavage and glycosylation in the Golgi [15, 16] prior to trafficking to cell-cell junctions [17, 18]. SPCA1 dysfunction particularly affects calcium-sensitive cadherins needed to build desmosomes, which are abundant in the epidermis, likely explaining HHD’s skin-focused phenotype. Severing of cadherin-based junctions between keratinocytes leads to epidermal splitting, called acantholysis, the pathologic hallmark of HHD [19, 20].

While its histologic features are well-established, HHD treatment is based mostly on case reports and expert opinion. Despite linkage of *ATP2C1* variants to HHD over two decades ago, there is no FDA-approved therapy [3, 21, 22]. Typical HHD treatments like corticosteroids or antimicrobials focus on secondary inflammation or infection, while rational therapeutics addressing its root cause remain elusive [4]. Unfortunately, the published murine model of *Atp2c1* inactivation did not exhibit skin blistering [23], impeding pre-clinical studies. Motivated by the limited understanding of HHD pathogenesis and the lack of proven treatments or a suitable animal model, we built human cellular and tissue models of HHD to discover its molecular drivers and identified viable drug repurposing opportunities [24].

## RESULTS

### SPCA1 deficiency weakens intercellular adhesion in human keratinocytes and organotypic epidermis

Since its pathology is localized to the epidermis, we engineered renewable and scalable models of HHD using TERT-immortalized human epidermal keratinocytes (THEKs; parent line N/TERT-2G [25]); these cells remain able to differentiate into a mature epidermis in organotypic cultures within 2 weeks [26, 27]. We used CRISPR/Cas9 in THEKs [28] to target indel mutations to an early exon in *ATP2C1* (or a pseudogene as a control), generating multiple heterozygous (HET) and homozygous knockout (KO) monoclonal lines. We confirmed SPCA1 protein depletion by immunoblotting (**Fig. 1A**).

**Figure 1:**
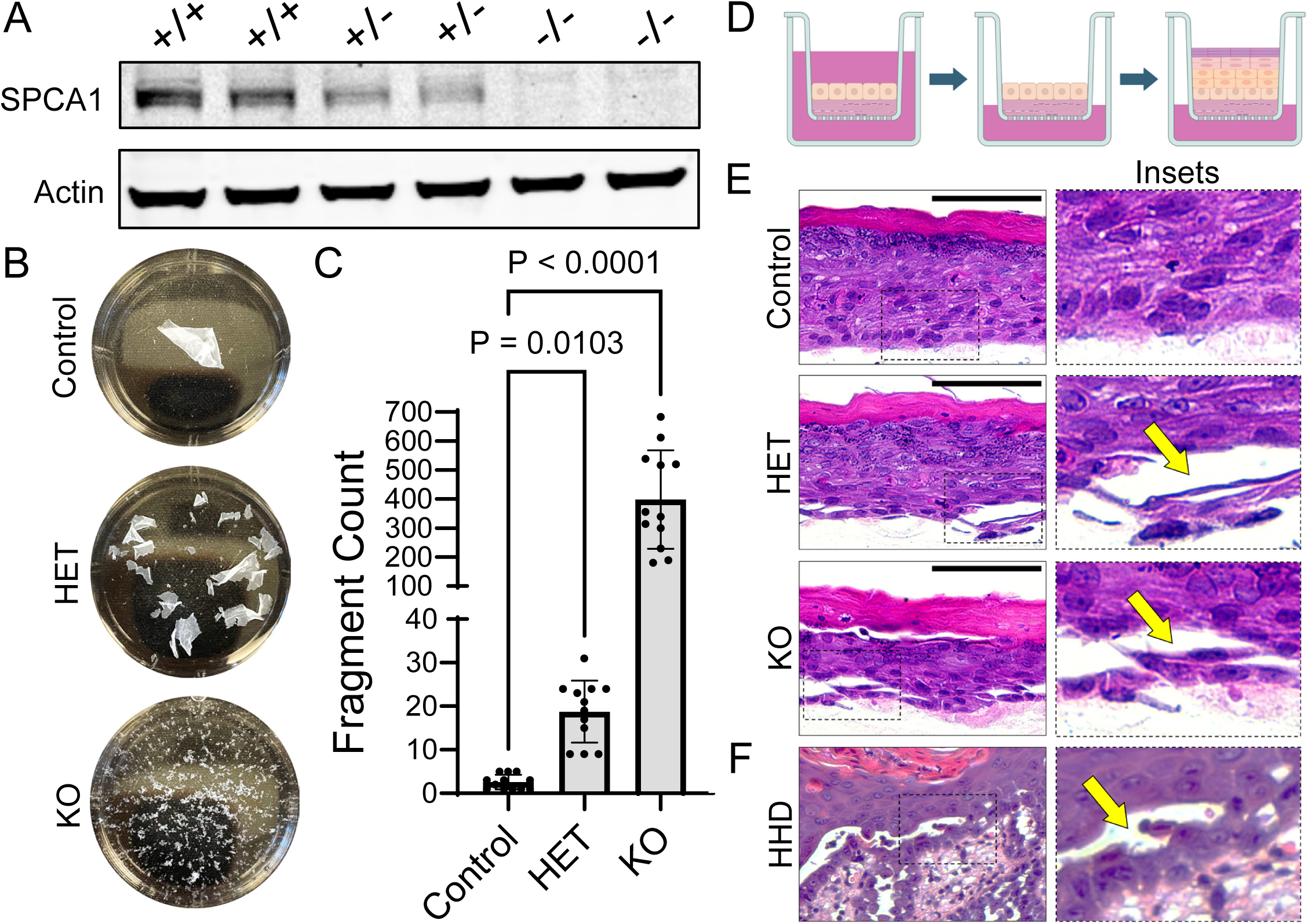
SPCA1 deficiency weakens intercellular adhesion in human keratinocytes and organotypic epidermis. (**A**) Immunoblot of SPCA1 in lysates from *ATP2C1* control (+/+), heterozygous (HET, +/-). and homozygous knockout (KO, -/-) TERT-immortalized human epidermal keratinocytes (THEKs); actin serves as a loading control. (**B**) Representative images show control, HET, and KO monolayers imaged in 35-mm wells after simultaneous application of equal mechanical stress. (**C**) Bar graph shows epithelial fragment counts (mean±SD) from mechanical dissociation assays (individual data points plotted for N=12 replicates; P-values from Welch’s ANOVA with Dunnett’s multiple comparison correction). (**D**) Schematic of organotypic epidermal culture model with keratinocytes seeded on collagen rafts, then grown at an air-liquid interface to induce stratification. (**E**) SPCA1-deficient organotypic epidermis displays acantholysis (arrows) mostly within the lower epidermal layers (insets magnified at right; bar=100 µm). (**F**) H&E-stained tissue section from a skin biopsy of a HHD lesion shows similar acantholysis (arrow, inset magnified at right).

To determine if SPCA1-deficient keratinocytes replicated the HHD phenotype, weakening of cell-cell adhesion, we deployed a mechanical dissociation assay validated to quantify desmosome strength [29]. Keratinocyte monolayers grown from HET and KO lines displayed increased fragmentation compared to controls (**Fig. 1B-C**). To assess their tissue-level phenotype, we generated epidermal cultures by growing THEKs at an air-liquid interface for 8 days (**Fig. 1D**). SPCA1-deficient organotypic epidermis displayed spontaneous splitting between keratinocytes, replicating the diagnostic feature of HHD (**Fig. 1E-F, arrows**).

### Depletion of SPCA1 impairs expression and localization of desmosomal proteins

To determine how SPCA1 deficiency weakens intercellular adhesion, we assessed expression and localization of cell-cell junction components. Immunoblotting of cell lysates demonstrated reduced desmosomal cadherins (desmogleins, DSG) in HET and KO keratinocytes (**Fig. 2A-B**). Immunostaining revealed that DSG3 and desmosomal plaque proteins (plakoglobin, desmoplakin) were less concentrated at intercellular junctions in HET and KO cells (**Fig. 2C-D**). Together, these data indicate that SPCA1 deficiency compromises both the level of cadherin proteins and their stability in cell-cell junctions, consistent with the fragility of HET and KO cell sheets (**Fig. 1B**).

**Figure 2:**
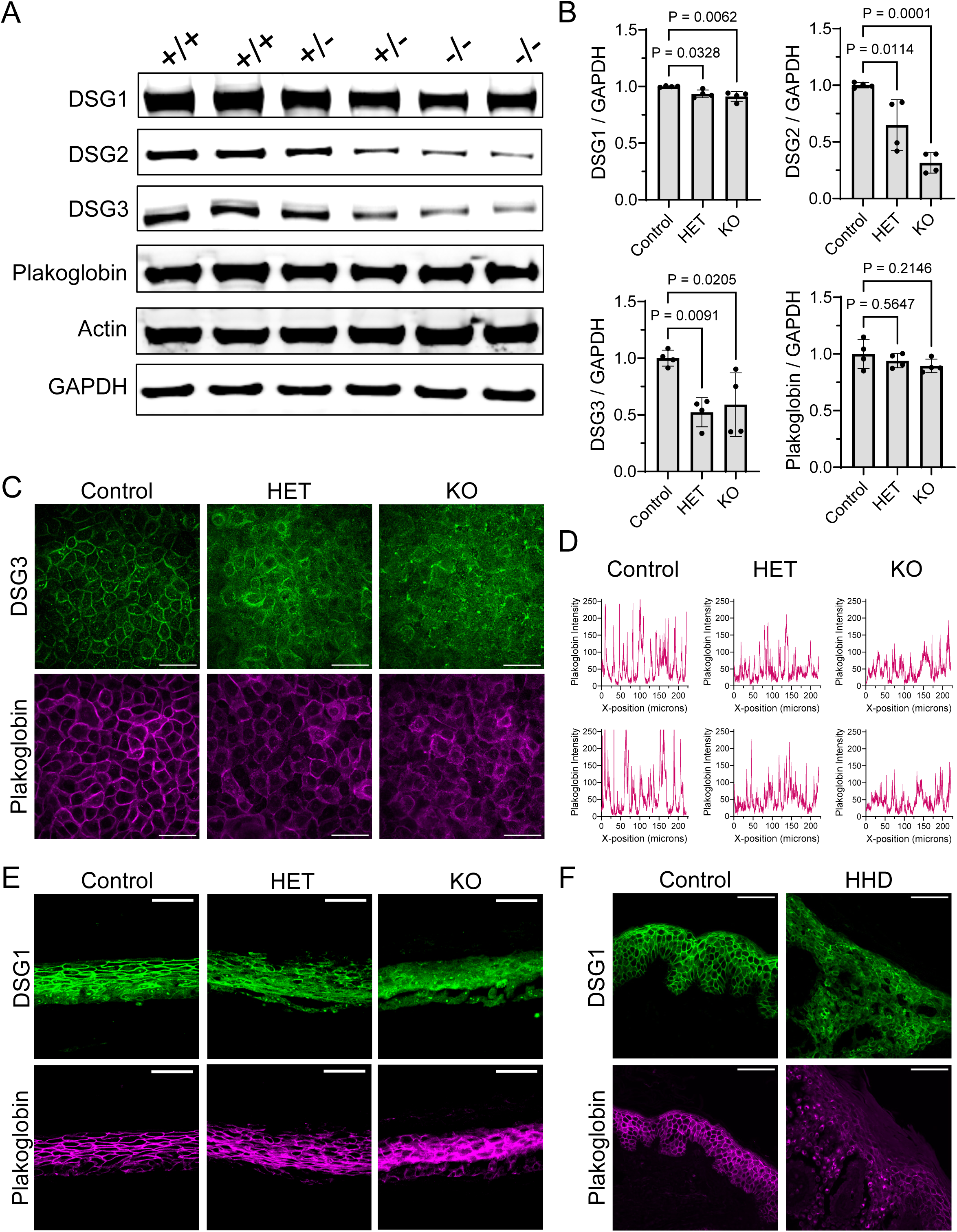
Depletion of SPCA1 impairs expression and localization of desmosomal proteins. (**A**) Immunoblot of lysates from 2 independent THEK lines for each genotype: control (+/+), HET (+/-), or KO (-/-); cells were exposed to 1.3 mM calcium for 4 hr prior to lysis. (**B**) Protein quantification (relative to GAPDH) showed decreased desmosomal cadherins (DSG1, DSG2, DSG3); bar graph shows mean±SD with individual data points plotted for N=4 experiments; P-values from 1-way ANOVA with Dunnett’s multiple comparison correction. (**C**) Immunostaining of DSG3 (green) and plakoglobin (magenta) in control, HET, and KO THEKs treated with 1.3 mM calcium for 4 hr prior to fixation in methanol (bar=50 µm); images representative of 2 independent lines per genotype. (**D**) Graphs depict plakoglobin fluorescence intensity at each pixel along a line-scan across representative confocal images from 2 cell lines per genotype; the largest peaks occur at well-formed cell-cell junctions. (**E**) Tissue cross-sections of organotypic cultures of HET and KO THEKs (bar=50 µm) and (**F**) HHD skin biopsies were immunostained for DSG1 (green) and plakoglobin (magenta), revealing reduced localization to intercellular borders compared to controls (bar=50 µm); images representative of N=5 organotypic tissues per genotype or N=5 donor biopsies per group.

We found similar results in SPCA1-deficient organotypic epidermis. Immunostaining of fixed tissue cross-sections demonstrated decreased DSG1 and plakoglobin at cell-cell borders (**Fig. 2E**). These results from our HHD model were substantiated by immunostaining HHD patient skin biopsies, which showed similar disruption of DSG1 and plakoglobin (**Fig. 2F**). Consistent with prior results from HHD biopsies [30–33], desmosomal proteins re-localized to the cytoplasm and peri-nuclear area, suggesting they may be internalized from the plasma membrane and/or stalled within secretory compartments.

### Chemical inhibition of SPCA1 disrupts intercellular junctions in primary human keratinocytes and organotypic epidermis

To complement this immortalized cellular model and validate the effects of genetically targeting *ATP2C1*, we optimized a second HHD model using chemical inhibition of SPCA1 in normal human epidermal keratinocyte (NHEKs). Previously published work identified 1,3-thiazole derivatives as specific SPCA1 inhibitors [34]. We tested two of these drugs (here termed compounds 3 and 5); both SPCA1 inhibitors decreased desmoglein protein levels in immunoblots of NHEK lysates (**Fig. 3A-B**) while plakoglobin was less affected. Immunostaining of inhibitor-treated NHEK monolayers revealed disrupted desmosomal proteins DSG3 and plakoglobin, similar to KO cells (**Fig. 3C-D**). Consistent with this impairment of junctional organization, NHEK monolayers treated with either SPCA1 inhibitor exhibited marked fragmentation in mechanical dissociation assays (**Fig. 3E**).

**Figure 3:**
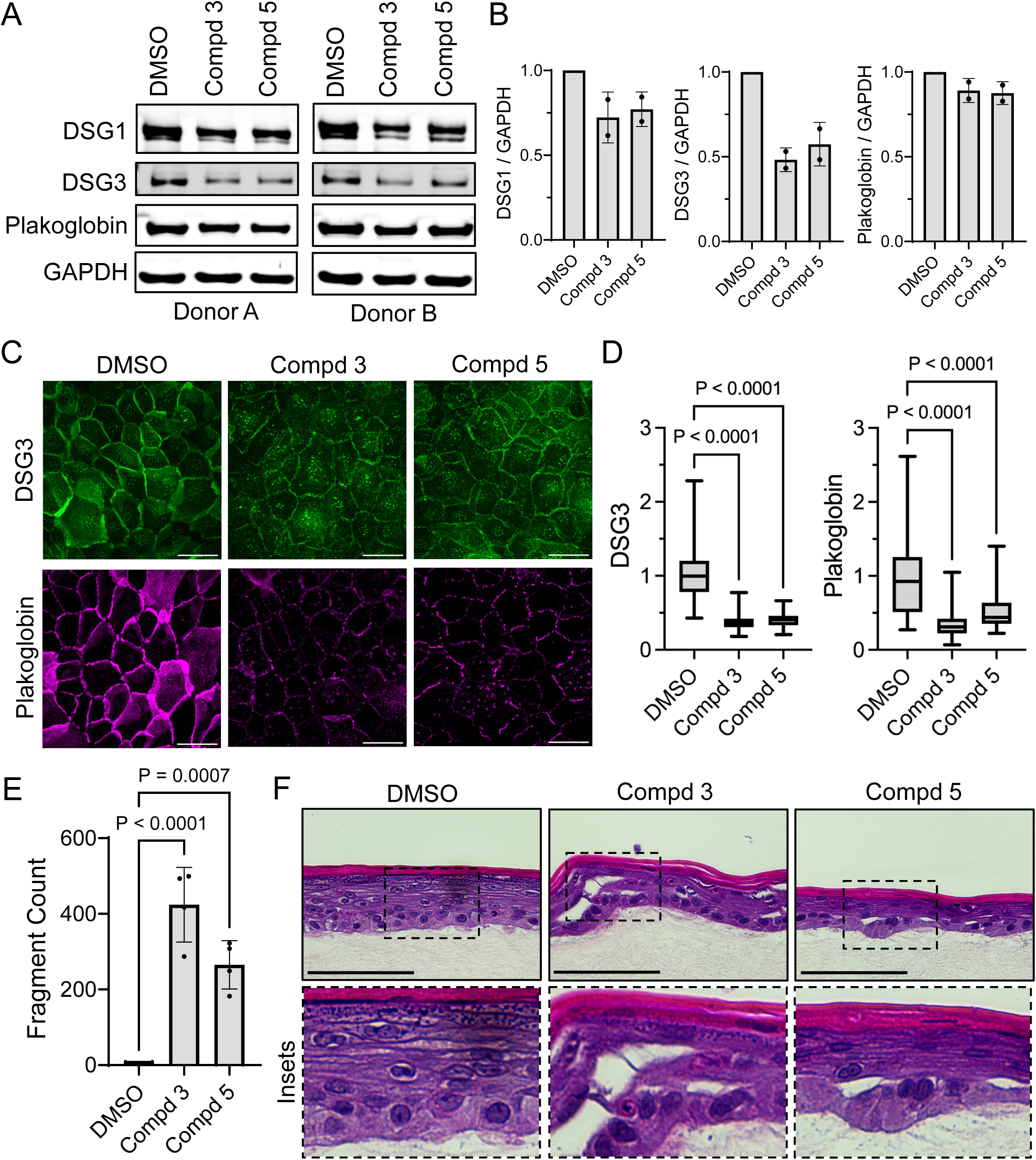
Chemical inhibition of SPCA1 disrupts intercellular junctions in primary human keratinocytes and organotypic epidermis. (**A**) Immunoblot of lysates from normal human epidermal keratinocytes (NHEKs) from 2 donors treated with 1.3 mM calcium along with DMSO or SPCA1 inhibitors, compound 3 (Compd 3) or 5 (Compd 5) at 10 ug/mL for 24 hr. (**B**) Protein quantification (relative to GAPDH) showed decreased desmosomal cadherins (DSG1, DSG3) with less effect on plakoglobin; bar graph shows mean±SD with data points plotted for N=2 keratinocyte donors. (**C**) Immunostaining of DSG3 (green) and plakoglobin (magenta) in NHEKs treated with 1.3 mM calcium plus DMSO vs. 10 ug/mL Compd 3 or 5 for 24 hr (bar=50 µm); images representative of N=3 NHEK donors. (**D**) Fluorescence intensity quantification showed a decrease in DSG3 and plakoglobin in NHEKs after 24 hr treatment with Compd 3 and 5 (data shown as a box plot of 25th–75th percentile with line at median; N≥57 images pooled from 3 donors; control mean normalized to 1; P-values from 1-way ANOVA with Dunnett’s multiple comparison correction). (**E**) Bar graph shows epithelial fragment counts (mean±SD) from mechanical dissociation assays; individual data points plotted for N=4 replicates; P-values from 1-way ANOVA with Dunnett’s multiple comparison correction. (**F**) H&E images of control THEK organotypic cultures treated with 40 ug/mL Compd 3 or 5 for 48 h; acantholysis was induced by both SPCA1 inhibitors (bar=100 µm; insets magnified below).

In mature organotypic epidermis treated with SPCA1 inhibitors for 48 hr, histologic analysis revealed acantholysis, particularly between the basal and first suprabasal layer (**Fig. 3F**), which is a common area of tissue splitting in HHD biopsies (**Fig. 1F**). Together these findings confirm the essential role of SPCA1 in the formation and function of desmosomes and corroborate the findings from our gene-edited THEK model. Further, our data support use of 1,3-thiazole derivatives to acutely block SPCA1, which replicated the key pathologic feature of HHD and offers an additional model for studying acantholytic disorders.

### RNA sequencing identifies actin cytoskeletal dysregulation in SPCA1-deficient cells

To identify pathogenic drivers mediating disruption of intercellular adhesion in our HHD model in a less biased manner, we performed triplicate bulk RNA sequencing (RNAseq) of 2 control and 2 HET keratinocyte lines. Differential expression analysis identified up- and down-regulated genes in HET cells (**Fig. 4A**). We utilized gene ontology (GO) to categorize transcripts that were significantly increased in HET keratinocytes, which mimic the SPCA1 haplo-insufficiency of HHD (**Fig. 4B**). The most up-regulated gene subsets had logical relevance to HHD, including genes associated with the actin cytoskeleton, cell-cell junctions, myosin II binding, cadherin binding, and GTPase activity. The implication of actin dysregulation in our model is consistent with *in vitro* analysis of HHD patient keratinocytes [35], electron microscopic studies of HHD skin lesions [33, 36], and recent RNAseq from HHD biopsies [37].

**Figure 4:**
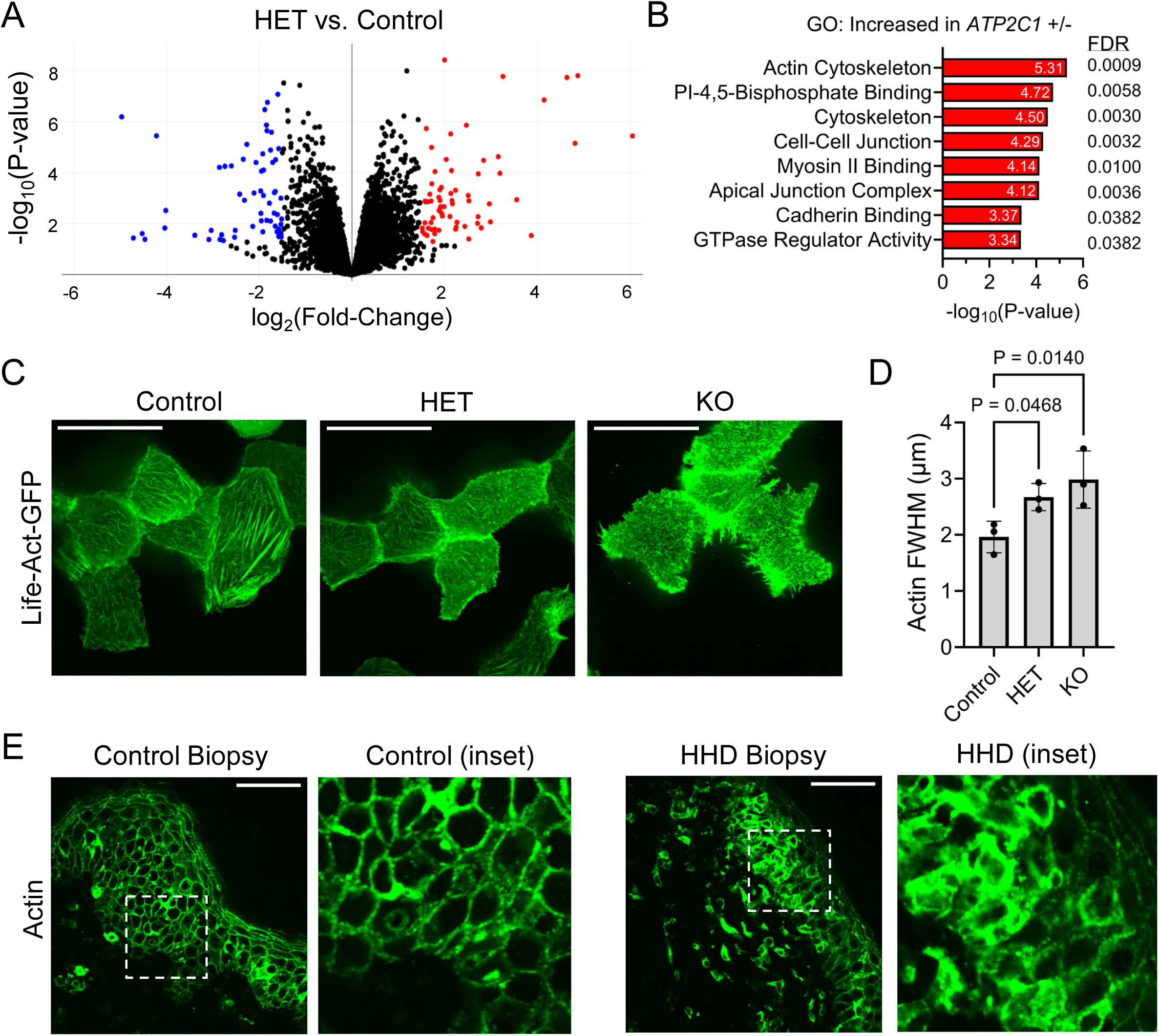
RNA sequencing identifies actin cytoskeletal dysregulation in SPCA1-deficient cells. (**A**) Volcano plot of bulk RNA sequencing (RNAseq) results from 2 HET vs. 2 control keratinocyte lines shows log_2_ of the fold-changes of genes significantly down- (blue) or up-regulated (red) with a cutoff of 0.05 for the adjusted P-value (plotted as -log_10_). (**B**) Gene ontology (GO) analysis of transcripts differentially expressed in 2 HET vs. 2 control keratinocyte lines (false discovery rate [FDR] and adjusted P-value both <0.05) revealed up-regulation of genes associated with the actin cytoskeleton, cell-cell junctions, myosin II binding, cadherin binding, and GTPase activity. (**C**) Confocal microscopy images (representative of N=3 experiments) of control, HET, and KO cells transduced with LifeAct-GFP reveal altered actin organization in SPCA1-deficient cells (bar=50 µm). (**D**) Cortical actin organization was quantified by measuring the full-width half-maximum (FWHM) of phalloidin intensity across intercellular borders; ≥109 borders measured per genotype; means plotted from N=3 experiments; P-values from 1-way ANOVA with Dunnett’s multiple comparison correction. (**E**) Immunostaining of control skin vs. HHD biopsies (representative of N=5 per group) reveals cortical actin disorganization in lesional areas (bar=50 µm; insets magnified at right).

Supporting our GO analysis pointing to actomyosin dysregulation, we found SPCA1-deficient keratinocytes exhibited marked differences in actin organization. Transduction of control, HET, and KO cells with LifeAct-GFP, a fluorescent peptide that allows real-time actin imaging, revealed that SPCA1-deficient cells displayed fewer stress fibers (**Fig. 4C**). In cells fixed and stained with phalloidin, the distribution of actin filaments at cell-cell junctions was widened in HET and KO lines (**Fig. 4D**), reflecting altered cortical cytoskeletal architecture. Likewise, HHD biopsies demonstrated cortical actin disruption near and within acantholytic areas (**Fig. 4E**). In agreement with prior observations [36], dissolution of cortical actin and adjacent cell-cell junctions is essential for acantholysis and suggests actin modulation may offer a therapeutic opportunity in HHD.

### Rho activation increases myosin light chain phosphorylation, disrupts intercellular junctions, and induces acantholysis

Given our RNAseq pointed to cytoskeletal regulators, including Rho family GTPases and myosin 2 (**Fig. 4B**), we hypothesized that RhoA, a major actin regulator in epidermis [38, 39], and its downstream targets could disrupt cortical actin and mediate contractile forces that drive intercellular junction severing in HHD. To test this, we treated control THEKs with calpeptin (CN01), which activates RhoA; CN01 dramatically weakened intercellular adhesion in keratinocyte monolayers, producing innumerable small fragments (**Fig. 5A**). We next tested whether Rho activation was sufficient to drive acantholysis by treating mature control organotypic cell cultures with CN01 for 48 hr. H&E staining revealed that CN01 induced splitting of keratinocyte junctions, particularly between the basal and suprabasal layers (**Fig. 5B**), mimicking the acantholysis we saw in SPCA1-deficient and -inhibited organotypic cultures as well as HHD biopsies (**Figs. 1E-F** and **3F**).

**Figure 5:**
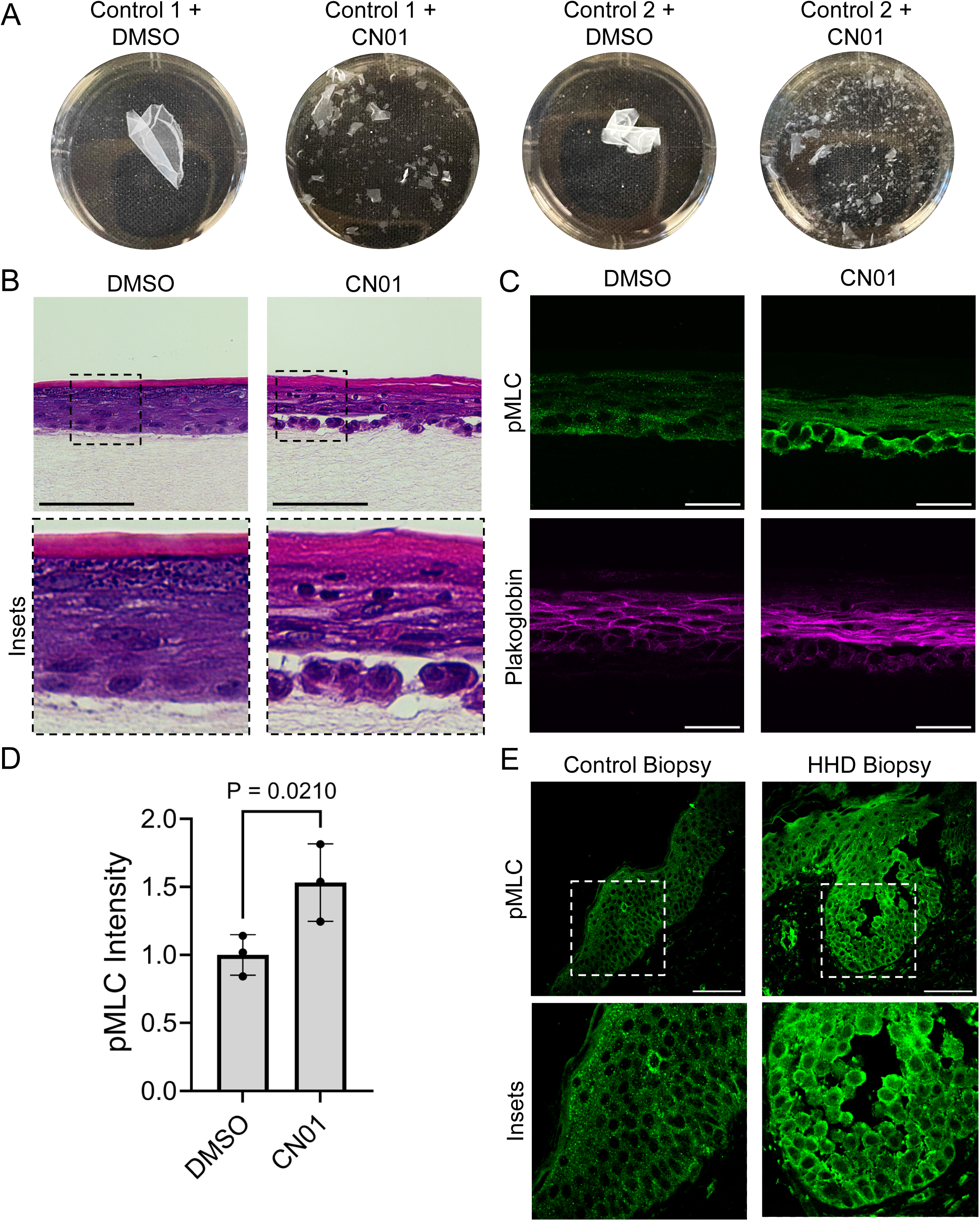
Rho activation increases myosin light chain phosphorylation, disrupts intercellular junctions, and induces acantholysis. (**A**) Representative images show control THEK monolayers treated with vehicle (DMSO) or Rho activator (CN01, 1 unit/mL); fragments were imaged in 35-mm wells after simultaneous application of equal mechanical stress. (**B**) H&E-stained tissue cross-sections of control organotypic cultures treated with DMSO or CN01 (0.5 units/mL) for 48 h; CN01 treatment induced acantholysis (bar=100 µm; insets magnified below); images representative of N=3 experiments. (**C**) Immunostaining of tissue cross-sections from organotypic cultures treated with DMSO or CN01 (0.5 unit/mL) showed increased pMLC (green) and disruption of plakoglobin (magenta) within acantholytic areas. (**D**) pMLC immunostaining quantification; bar graph displays mean±SD from ≥23 images per condition with average intensities plotted from N=3 experiments; mean of controls normalized to 1; P-value from 2-tailed paired t-test. (**E**) Immunostaining of tissue cross-sections from control skin or HHD biopsies highlights increased pMLC near and within acantholytic regions (insets magnified below); images representative of N=5 biopsies per group.

To understand how Rho disrupts cell-cell adhesion, we assessed its direct target, Rho-associated protein kinase (ROCK), which phosphorylates and activates myosin light chain (MLC) to exert tension on actin filaments. Immunostaining demonstrated a clear increase in phosphorylated myosin light chain 2 (pMLC) in CN01-treated organotypic epidermal cultures (**Fig. 5C-D**). These results are further supported by immunostaining of HHD biopsies, which showed markedly enhanced pMLC within and at the periphery of acantholytic foci (**Fig. 5E**). These findings led us to test if blunting ROCK, the upstream MLC activator (**Fig. 6D**), could rescue cohesion of SPCA1-deficient keratinocyte sheets.

**Figure 6:**
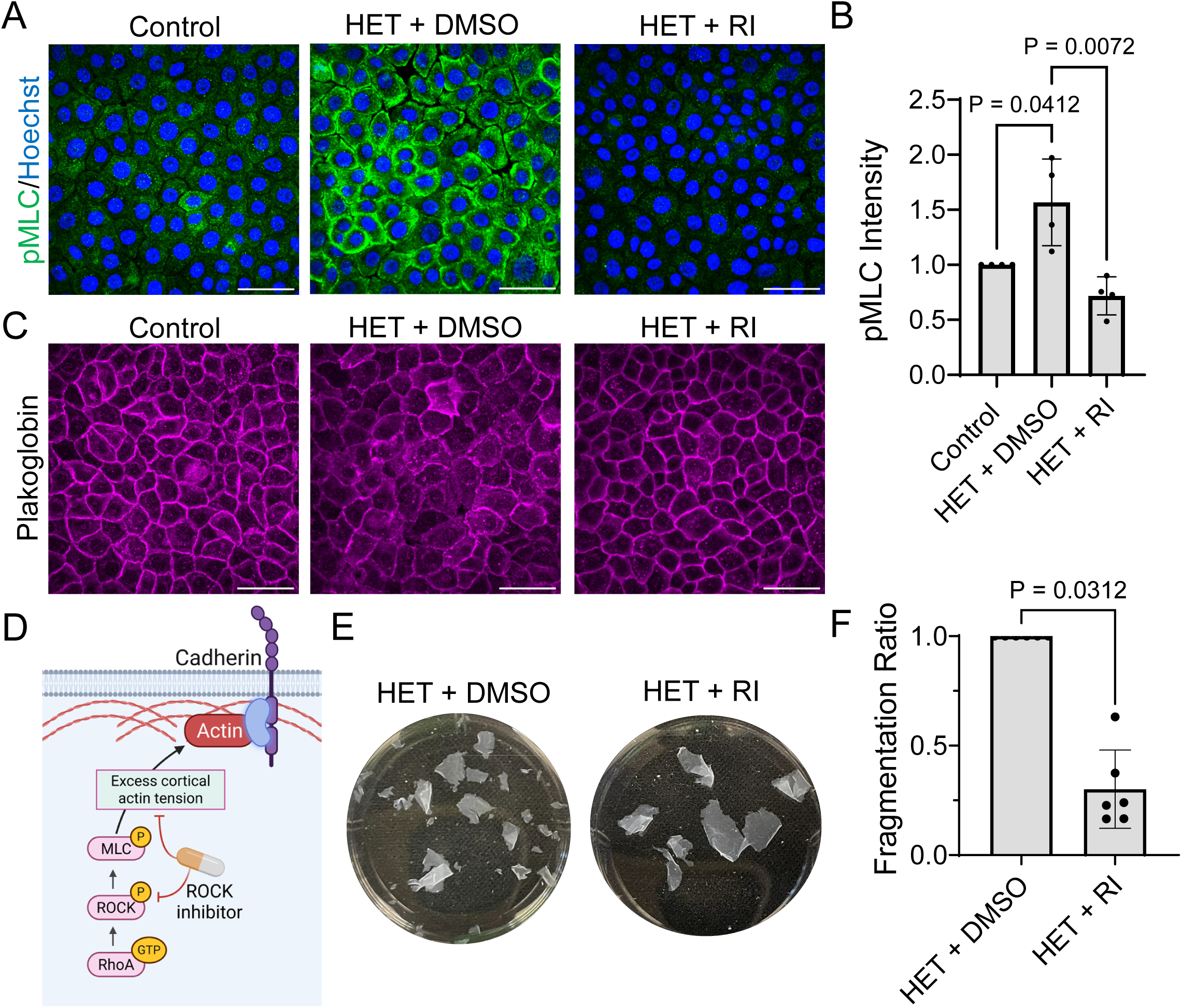
ROCK inhibition bolsters cell-cell junctions in SPCA1-deficient keratinocytes. (**A**) Immunostaining of phosphorylated myosin light chain 2 (pMLC, green; Hoechst, blue) in control and HET keratinocytes exposed to 1.3 mM calcium for 4 hr prior to fixation (bar=50 µm); cells were treated with DMSO or a ROCK inhibitor (RI; Y27632, 10 µM); images representative of N=2 cell lines per genotype. (**B**) Quantification of pMLC fluorescence intensity from N=4 replicates; P-values from 1-way ANOVA with Dunnett’s correction for multiple comparisons. (**C**) Representative images from N=4 experiments of plakoglobin immunostaining in cells treated as in (A) showing cell-cell junction compromise in HET cells and rescue by ROCK inhibition (RI). (**D**) Diagram of RhoA activation of ROCK and MLC, which results in excess cortical actin tension at cell-cell junctions. (**E**) Representative images show keratinocyte monolayers treated with vehicle (DMSO) or a ROCK inhibitor (RI; Y27632, 4 µM); fragments were imaged in 35-mm wells after simultaneous application of equal mechanical stress. (**F**) Bar graph shows the ratio of epithelial fragment counts (mean ± SD) from mechanical dissociation assays using ROCK-inhibitor (RI) vs. DMSO treatment; points plotted for N=6 replicates pooled from 2 HET lines; P-value from Wilcoxon rank-sum test.

### ROCK inhibition bolsters cell-cell junctions in SPCA1-deficient keratinocytes

Given the increased pMLC in blistered areas of HHD biopsies, we examined whether this regulator of cortical actin tension was activated in our HHD models. Compared to control cells, immunostaining of HET keratinocytes revealed higher pMLC fluorescence intensity, including at cell-cell borders (**Fig. 6A-B**). Thus, we reasoned that dampening MLC activation using a ROCK inhibitor might blunt its effect on cell-cell junctions. Accordingly, treating HET keratinocytes with a ROCK inhibitor (Y27632, 10 µM) decreased pMLC (**Fig. 6B**), which correlated with an increase in junctional localization of plakoglobin (**Fig. 6C**), suggesting ROCK inhibition can bolster cell-cell junctions despite SPCA1 deficiency.

We next assessed if a ROCK inhibitor could rescue the intercellular adhesive strength of HET keratinocytes. In cell sheets grown from either HET line, we found a reduction in the number of fragments after mechanical dissociation when they were pre-treated with Y27632, though the fragmentation of cell sheets was not fully halted (**Fig. 6E-F**). These results suggest that ROCK inhibitors could fortify junctions and potentially reduce blistering in HHD. Since their effect was moderate, we were motivated to investigate additional interventions to more robustly rescue intercellular adhesion.

### SPCA1-deficient cells exhibit ERK hyperactivation and benefit from MEK inhibition

Like HHD, Darier disease (DD) is caused by deficiency of a calcium pump, SERCA2. In our published *in vitro* model of DD, we found that increased activation of the mitogen-activated protein (MAP) kinase ERK weakened cell-cell adhesion and identified MEK inhibition as an effective therapy [40]. These results were corroborated in a model of Grover disease, an idiopathic acantholytic disorder that we also linked to ERK hyperactivation [41]. Interestingly, RNAseq of *ATP2C1* HET keratinocytes identified differential expression of regulators of cell-cell junctions and cadherin-mediated adhesion (**Fig. 4B**). This included dysregulation of the epidermal growth factor (EGF) receptor, which activates downstream signaling via MEK and ERK that have been linked to acantholytic blistering diseases [40–44].

To determine if HET and KO keratinocytes displayed differences in ERK signaling, we utilized a genetically encoded biosensor. The ERK kinase translocation reporter linked to a green fluorophore (ERK-KTR-mClover)[45] remains in the cytoplasm when active and localizes to the nucleus when inactive (**Fig. 7A**). After treating THEKs with 1.3 mM calcium for 4 hr, HET and KO keratinocytes displayed more cytoplasmic fluorescence, indicating ERK activation, while control cells displayed more nuclear signal (**Fig. 7B**). As in prior work [41], we quantified the cytoplasmic/nuclear fluorescence intensity ratio as an ERK activity index, which confirmed that HET and KO cells had a significant increase in ERK activity (**Fig. 7C**). These results suggested dampening ERK with an upstream MEK inhibitor could be therapeutic in our HHD model.

**Figure 7:**
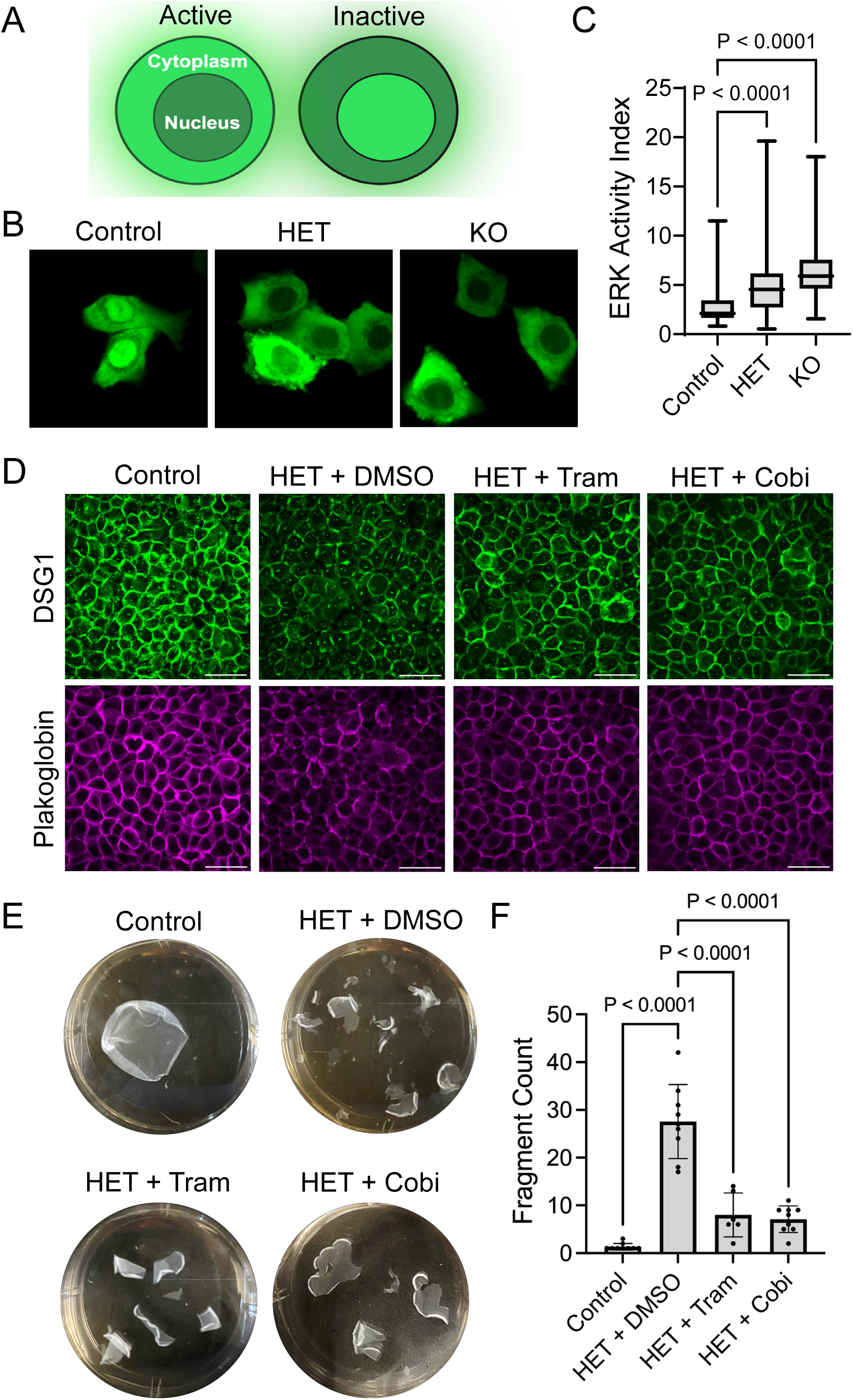
SPCA1-deficient cells exhibit ERK hyperactivation and benefit from MEK inhibition. (**A**) Schematic of an ERK biosensor (ERK-KTR-mClover), which localizes the mClover fluorophore to the cytoplasm when the kinase is active and the nucleus when inactive. (**B**) Representative images of control, HET, and KO keratinocytes transduced with the ERK biosensor; cells were treated with 1.3 mM calcium for 4 hr prior to imaging. (**C**) The ERK activity index was calculated as the cytoplasmic-to-nuclear ratio of green fluorescence intensity; data shown as a box plot of 25th–75th percentile with line at median for values pooled from N≥600 cells per group from 2 cell lines per genotype from 2 replicate experiments; P-values from 1-way ANOVA with Dunnett’s multiple comparison correction. (**D**) Representative images of immunostaining of DSG1 (green) and plakoglobin (magenta) in THEKs after exposure to 1.3 mM calcium for 4 hr; HET cells were treated with DMSO or a MEK inhibitor (cobimetinib [Cobi] or trametinib [Tram], both 1 µM); images representative of 2 cell lines per genotype, pooled from N=3 experiments. (**E**) Representative images show THEK monolayers treated with DMSO or a MEK inhibitor (Cobi or Tram, both 1 µM) plus 1.3 mM calcium for 18 hr; fragments were imaged in 35-mm wells after simultaneous application of equal mechanical stress. (**F**) Bar graph shows epithelial fragment counts (mean±SD) from mechanical dissociation assays; individual data points plotted for N=6 or 9 replicates pooled from 3 experiments; P-values from 1-way ANOVA with Dunnett’s multiple comparison correction.

We treated HET keratinocytes with MEK inhibitors cobimetinib (Cobi, 1 µM) or trametinib (Tram, 1 µM) for 18 hr. DSG1 and plakoglobin immunostaining demonstrated that either MEK inhibitor increased localization of these desmosomal proteins to intercellular junctions in HET monolayers (**Fig. 7D**). This translated into an increase in adhesive strength in mechanical dissociation assays; HET monolayers treated with either MEK inhibitor had a lower fragment count than those treated with DMSO (**Fig. 7E-F**), While the inhibitors did not achieve a complete rescue of integrity, these drugs, which are already FDA-approved for cancer [46] and are used off-label for dermatological diseases driven by MAP kinase signaling [47–49], might be re-purposed to halt or prevent blistering in an orphan disorder like HHD.

### MEK and ROCK inhibitors demonstrate synergistic therapeutic effects in HHD model

Given the partial rescue of HET keratinocyte sheet integrity achieved by ROCK or MEK inhibitors, we reasoned they might have synergistic effects. We treated SPCA1-deficient monolayers with a ROCK inhibitor (Y27632, 4 µM) plus a MEK inhibitor (Cobi, 1 µM) for 18 hr before applying mechanical stress. HET keratinocyte sheets treated with the dual inhibitors displayed markedly increased plakoglobin at intercellular junctions compared to DMSO-treated cells (**Fig. 8A-B**). Excitingly, this enhancement of desmosomal proteins at cell-cell borders translated into an increase in monolayer integrity. In fact, HET cell sheets exhibited comparable strength to controls, fully resisting fragmentation in mechanical dissociation assays (**Fig. 8C-D**).

**Figure 8:**
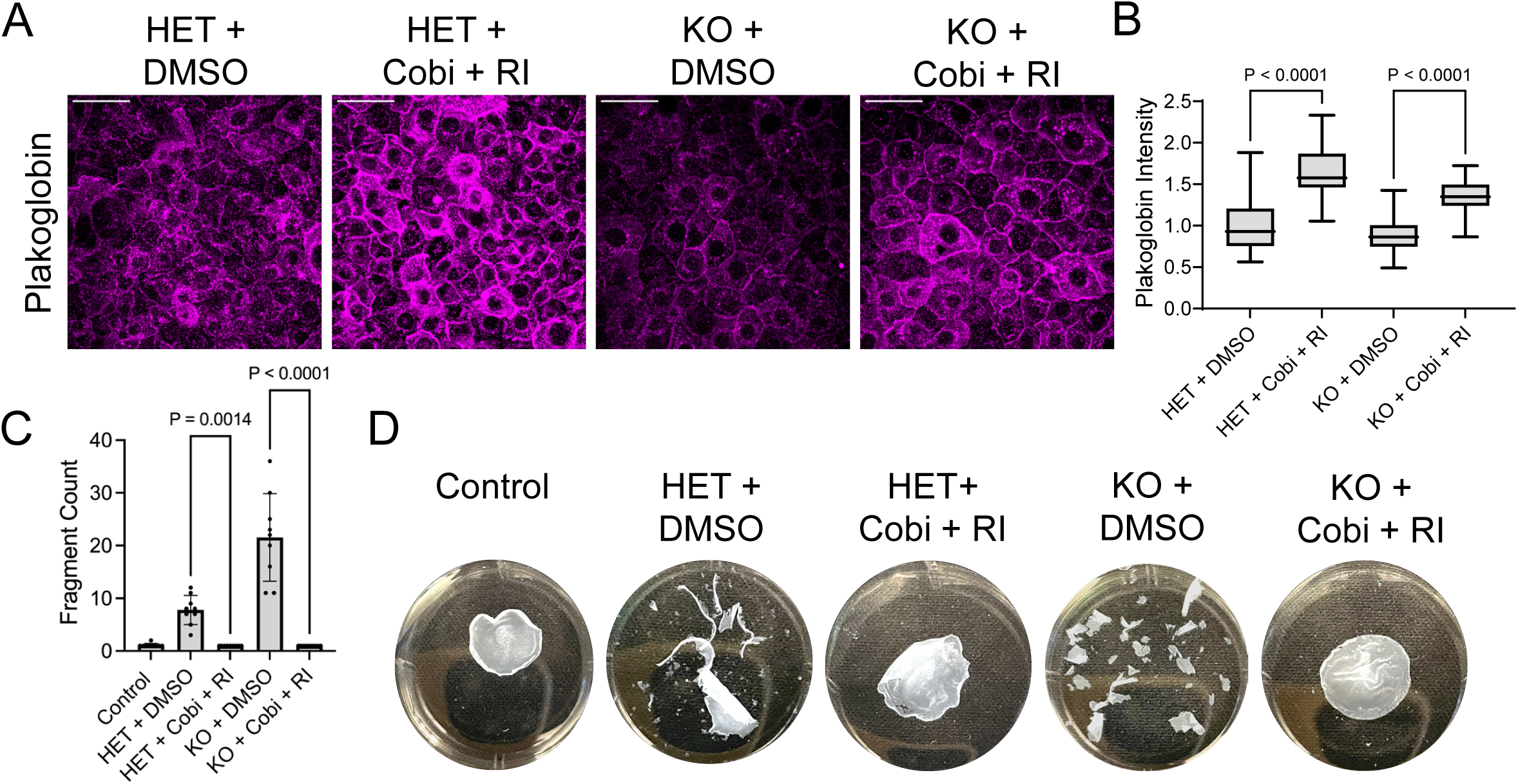
MEK and ROCK inhibitors demonstrate synergistic therapeutic effects in HHD model. (**A**) Representative images of plakoglobin immunostaining in HET and KO cell lines exposed to 1.3 mM calcium for 4 hr prior to fixation (bar=50 µm); cells were treated with vehicle (DMSO) or a MEK inhibitor (cobimetinib [Cobi], 1 µM) plus a ROCK inhibitor (Y27632 [RI], 4 µM). (**B**) Quantification of plakoglobin fluorescence intensity; data shown as a box plot of 25th–75th percentile with line at median for values pooled from N≥80 images per group from 2 cell lines per genotype; P-values from 1-way ANOVA with Bonferroni’s multiple comparison correction. (**C**) Bar graph shows epithelial fragment counts (mean±SD) with data plotted for N=9 replicates; P-values from 1-way ANOVA with Bonferroni’s multiple comparison correction. (**D**) Representative images of THEK monolayers treated with DMSO or a MEK inhibitor (Cobi, 1 µM) plus a ROCK inhibitor (Y27632, 4 µM); fragments were imaged in 35-mm wells after simultaneous application of equal mechanical stress.

Encouraged by these findings, we tested the effect of the dual inhibitors on homozygous KO keratinocyte sheets completely lacking functional SPCA1, which had not been strengthened significantly by either ROCK or MEK inhibitors alone (not shown). Remarkably, treatment with both a ROCK and MEK inhibitor promoted plakoglobin localization to intercellular borders, even in KO keratinocytes, and restored their mechanical integrity to control levels.

## DISCUSSION

A significant factor limiting our understanding of HHD has been the lack of an optimal pre-clinical model. In 2007, Okunade *et al*. reported that *Atp2c1* heterozygous mice did not exhibit skin blistering and homozygous KO mice were not viable [23]. *Saccharomyces cerevisiae* have been used as a simpler model to confirm that HHD-linked variants in the yeast *ATP2C1* homolog *pmr1* impaired the calcium pump’s function and induced secretory compartment stress [50, 51]. However, single-celled yeast lack intercellular junctions, the major target of HHD pathology. Other published *in vitro* HHD models utilized short-lived primary keratinocytes from patients or transient SPCA1 knockdown in immortalized human cell lines that do not produce a fully differentiated tissue [35, 52–55]. While such models exhibit disruption of desmosomes and the cytoskeleton as in HHD, they present limitations in experiment length and replication due to primary keratinocyte lifespan or variable and transient SPCA1 depletion.

Our study overcomes several of these limitations by generating a renewable cell culture model for HHD utilizing CRISPR/Cas9 to stably ablate *ATP2C1* in N/TERT-immortalized human keratinocytes [56, 57], which remain capable of generating a differentiated epidermal tissue [25]. We isolated multiple homozygous knockout (KO) and heterozygous (HET) cell lines, which replicated the HHD phenotype. As monolayers, SPCA1-deficient cells exhibited weakened cohesion in mechanical assays, which allowed testing of drugs for therapeutic effects; in organotypic cultures, they exhibited spontaneous acantholysis, the diagnostic feature of HHD. We also deployed chemical inhibitors [34] to acutely block SPCA1 in primary human keratinocytes, generating a secondary approach for studying HHD. Combining these models with miniaturized skin-on-chip platforms [58] could further expand their efficiency and scalability for drug screening [24].

Studying rare disorders like HHD can have implications for additional diseases [59]. Acantholytic blistering disorders are defined by their common pathologic feature, severing of epidermal cell-cell junctions [19, 20]. These include: Darier disease (DD), a ‘sister disease’ of HHD caused by variants in *ATP2A2*, which encodes the endoplasmic reticulum calcium pump SERCA2 [60]; Grover disease, an idiopathic disorder recently linked to *acquired ATP2A2* mutations [61]; and pemphigus, mediated by auto-antibodies against desmosomal cadherins [62, 63]. All are hallmarked by painful skin breakdown and recurrent infections. Despite decades of research, no therapies directly targeting the intercellular adhesive machinery have advanced to clinical trials. Our transcriptomic analysis identified two candidate HHD drivers that we validated in functional *in vitro* assays and HHD biopsies. Suppressing MAP kinase signaling via ERK bolstered desmosomal cadherins and inhibiting ROCK dampened myosin activation; together, these strategies rescued the integrity of SPCA1-deficient keratinocyte sheets. Importantly, both kinases are targeted by drugs FDA-approved for other diseases [46, 64–66].

Transcriptomic analysis of HET cells led us to hypothesize that weakening of cell-cell adhesion was driven by EGF receptor signaling, which activates RAS, RAF, and MAP kinases known to regulate desmosomes [67–69]. We validated this finding in HET cells using an ERK biosensor (**Fig. 7A-C**). EGF receptor signaling drives human squamous cell carcinomas [70], thus excess activation of this pathway may promote the age-related skin tumors seen in *Atp2c1* [23] and *Atp2a2* [71] HET mice, the latter showing markedly increased RAS [72]. Overactivity of this pathway may also explain a slight increase in keratinocyte carcinomas in DD patients [73]. Interestingly, Grover disease, which exhibits acantholysis similar to HHD, can be induced by hyperactive ERK as a paradoxical side effect of BRAF inhibitors [74–76], which have also been linked to keratinocyte neoplasms [77].

Our work shows that using MEK inhibitors to dampen ERK signaling strengthens cell-cell adhesion in SPCA1-deficient keratinocytes. This result is consistent with the therapeutic effects of MEK blockade seen in our model of DD [40], which informed treatment of a severely affected patient using a MEK inhibitor, trametinib [78]. While prior work linked acantholysis in pemphigus to MAP kinase activation [43, 79], most studies focused on blocking a different MAP kinase, p38 [80, 81], a strategy that was not viable in subsequent work [82] or in our HHD model (not shown). However, a recent study showed erlotinib, an EGF receptor inhibitor that suppresses downstream MEK and ERK, prevented blistering in a human explant model of pemphigus [42]. Indeed, EGF receptor modulation has emerged as a promising strategy for several epidermal differentiation disorders [83–85]. Our studies suggest that selective MEK or ERK inhibitors may offer a more targeted strategy to strengthen epidermal integrity in these orphan diseases.

Our results are consistent with recent *in vitro* studies identifying actin dysregulation in keratinocyte models of HHD [86, 87] and in bulk transcriptomic analysis of HHD biopsies [88]. Though the latter approach could not rule out a contribution of inflammatory and wound responses, our RNAseq identified keratinocyte-autonomous over-activation of actomyosin modulators, allowing us to identify a therapeutic effect of Rho kinase (ROCK) inhibition. ROCK, which is known to regulate cell-cell junction strength, epidermal stratification, and wound repair [38, 39, 89], phosphorylates the light chain of myosin motors (pMLC) to exert tension on cortical actin [90–92]. In our cellular and organotypic models of HHD, as well as in patient biopsies, we identified increased pMLC in acantholytic areas. We propose this reflects actomyosin tension that ‘pulls’ keratinocytes apart during acantholysis. RhoA and ROCK have been linked to the stability of desmosomes [93, 94]. While some reports suggest RhoA and ROCK inhibition would be therapeutic for pemphigus [95], others proposed that RhoA drives blistering [96, 97]. In our experiments, a RhoA activator induced robust tissue splitting; this was paralleled by increased pMLC near and within acantholytic areas of organotypic cultures and HHD biopsies (**Fig. 5**). Differences in dosing or timing of RhoA modulation or distinct responses to direct desmosome disruption by auto-antibodies in pemphigus may explain these discrepancies.

For orphan diseases like HHD, developing new therapies can prove challenging given the limited number of patients as well as the risk and expense of early-phase clinical studies; however, drug re-purposing is a viable approach for rare disorders [24, 98]. Drugs that are FDA-approved for other indications can be prescribed off-label, which has been done for acantholytic disorders [20]. While case reports suggest potential utility of anti-cytokine antibodies in HHD [99–102], these target inflammation that may be secondary to the skin barrier breach; in fact, their initial effects may not be sustained based on follow-up studies [103, 104]. Instead, our analysis points to inhibitors of MEK and ROCK, several of which are FDA-approved or under investigation for other indications, including dermatologic diseases. MEK inhibitors are a first-line therapy for BRAF-driven melanoma [105] and have been used off-label for vascular anomalies [47], while ROCK inhibitors may treat keloid scars [106–108]. MEK inhibitors have been compounded for topical treatment of cutaneous histiocytosis [109] and neurofibromatosis [110, 111] and ROCK inhibitor eyedrops are FDA-approved for glaucoma [66, 112].

External application of MEK or ROCK inhibitors is feasible, including for the drugs used in our studies [109, 113, 114], and would limit toxicity. Moreover, blistering diseases present a unique susceptibility to topical drug delivery given their breach of the skin barrier. Finally, beyond the specific advance from our studies identifying therapeutic strategies for HHD, our approach to HHD modeling that combined gene editing with organotypic epidermis provides a blueprint for replicating other rare epidermal disorders. Moreover, gene-edited epidermal disease models provide a renewable source of diseased tissue for pre-clinical vetting of emerging therapeutic technologies for topical gene replacement [115, 116] or correction of pathogenic variants using precise DNA editing tools such as CRISPR, Prime editors, and base editors [117], which were recently deployed in an epidermal differentiation disorder [118].

## METHODS

### Sex as a biological variable

Our studies used available human keratinocytes (primary and immortalized) from male subjects; however, we validated *in vitro* results in skin biopsies from both females and males, yielding similar findings.

### Reagents

Dimethyl sulfoxide (DMSO; Cat. #BP231-100) was from Fisher Scientific. Published and additional data from Dr. Sachiko Yamamoto-Hijikata (Kyorin Univ. School of Medicine, Mitaka, Tokyo) identified 1,3-thiazole derivatives, specifically [4-(dimethylamino)-N-[4-(4-methylphenyl)-1,3-thiazol-2-yl]benzamide] as specific inhibitors of SPCA1 [34]. These compounds are identified as compound 3 (Enamine Cat. #T5370096) and compound 5 (Life Chemicals Cat. #F0375-0126). MEK inhibitors include trametinib (Cell Signaling Cat. #62206) and cobimetinib (GDC-0973, SelleckChem Cat. #S8041). All chemicals were used as specified in figure legends.

Primary antibodies included: Rabbit anti-SPCA1 from Sigma (Cat. #HPA069684); rabbit phospho-ERK1/2 (D13.14.4E; Cat. #4370) and mouse anti-ERK1/2 (L34F12; Cat. #4696) and anti-β-Actin (8H10D10; Cat. #3700) from Cell Signaling Technologies; mouse anti-desmoglein 1 (Cat. #ab12077) from Abcam; mouse anti-desmoglein 2 (AH12.2; Cat. #sc-80663), mouse anti-desmoglein 3 (5G11; Cat. #sc-53487), mouse anti-plakoglobin (A-6; Cat. #sc-514115), mouse anti-β-Actin (C4; Cat. #sc-47778; immunoblotting 1:500), and mouse anti-GAPDH (0411; Cat. #sc-47724; immunoblotting 1:500) from Santa Cruz.

Secondary antibodies for fluorescent immunoblotting (1:10,000), IRDye 800CW goat anti-rabbit IgG (Cat. #926-32211) and IRDye 680RD goat anti-mouse IgG (Cat. #926-68070), are from LI-COR Biosciences. Secondary antibodies for immunostaining of cells and tissues (1:300) were from Thermo-Fisher: Goat anti-mouse IgG AlexaFluor-405 (Cat. #A31553), AlexaFluor-488 (Cat. #A11001), AlexaFluor-594 (Cat. #A11005), or AlexaFluor-633 (Cat. #A21050); goat anti-rabbit

IgG AlexaFluor-405 (Cat. #A31556), AlexaFluor-488 (Cat. #A11008), AlexaFluor-594 (Cat. #A11012), or AlexaFluor-633 (Cat. #A21070); goat anti-chicken IgY AlexaFluor-488 (Cat. #A-11039), AlexaFluor-594 (Cat. #A-11042), or AlexaFluor-633 (Cat. #A-21103). Hoechst 33342 is from Thermo-Fisher (Cat. #H1399; immunostaining 20 μg/ml).

### Cell culture

All cell lines were cultured at 37°C in an air-jacketed humidified incubator with 5% CO_2_.

Cells were cultured on sterile tissue culture-treated plates and passaged to remain sub-confluent using 0.25% Trypsin-EDTA (Thermo-Fisher, Cat. #15400054).

Normal human epidermal keratinocytes (NHEKs) were isolated from de-identified male neonatal foreskins by the Penn Skin Biology and Disease Resource-based Center. Cells were cultured in Medium 154 with 0.07 mM CaCl_2_ (Thermo-Fisher, Cat. #M154CF500) with 1x human keratinocyte growth supplement (Thermo-Fisher, Cat. #S0015) plus 1x gentamicin/amphotericin (Thermo-Fisher, Cat. #R01510).

hTERT-immortalized human epidermal keratinocytes (THEKs) from the original N/TERT-2G line [25] were grown in keratinocyte serum-free medium (KSFM) from Thermo-Fisher (Cat. #37010022) with 0.2 ng/mL human epidermal growth factor, 30 µg/mL bovine pituitary extract, plus 0.31 mM CaCl_2_, 100 U/mL penicillin, and 100 μg/mL streptomycin.

J2-3T3 immortalized murine fibroblasts (a gift from Dr. Kathleen Green, Northwestern University, Chicago, IL, USA) were grown in complete Dulbecco’s Modified Eagle Medium (DMEM) (Thermo-Fisher, Cat. #11965092) with 10% Hyclone FBS (Fisher Scientific, Cat. #SH3039603), 2 mM GlutaMAX (Thermo-Fisher, Cat. #35050061), plus 100 U/mL penicillin and 100 μg/mL streptomycin.

### CRISPR/Cas9 gene editing

CRISPR knockout (KO) keratinocytes were generated as described [56]. Single-guide RNAs (sgRNAs) were designed to target *ATP2C1* (sgRNA: GGAGCTGTCACCTTAGAACA) or the *TUBAP* pseudogene (sgRNA: GTATTCCGTGGGTGAACGGG) to generate control KO lines. A web tool was used for CRISPR strategy (https://portals.broadinstitute.org/gpp/public/analysis-tools/sgrna-design). Synthetic sgRNA target sequences were introduced into a cloning backbone, pSpCas9 (BB)-2A-GFP (PX458) (Addgene, Cat. #48138), then were cloned into competent *E. coli* (Thermo-Fisher, Cat. #C737303). Sanger sequencing was used to confirm proper insertion. Final plasmids were transfected into N/TERT-2G keratinocytes [25] using a TransfeX kit (ATCC, Cat. #ACS4005) with or without the JAK1/JAK2 inhibitor baricitinib (10 µg/mL). Single GFP-positive cells were selected and expanded. The presence of heterozygous or homozygous mutations in the targeted gene was confirmed via Sanger sequencing.

### RNA sequencing (RNAseq)

RNAseq libraries were prepared for transcriptomics analysis as described [119]. THEKs were grown to confluency in 6-well plates in KSFM, then differentiated in E-medium for 72 hours. RNA was isolated from THEKs utilizing RNeasy kit (Qiagen) according to the manufacturer’s instructions. NEBNext Poly(A) mRNA magnetic isolation module (New England Biolabs) was used to isolate mRNAs. NEBNext Ultra-Directional RNA library preparation kit was used to prepare RNAseq libraries for Illumina (New England Biolabs). Agilent BioAnalyzer 2100 was used to confirm library quality and the NEBNext Library Quant Kit for Illumina (New England Biolabs) was used to quantify. Sequencing was performed using the Illumina NextSeq500 platform utilizing a single-end, 75-base pair sequencing strategy.

RNAseq FASTQ files were processed using the cloud-based platform Elysium (https://maayanlab.cloud/cloudalignment/elysium.html) [120], which utilizes pseudoalignment to generate gene counts through the RNAseq quantification program Kallisto [121]. Kallisto assigns reads to compatible transcripts without performing base-level alignment, instead utilizing k-mer base matching to produce transcript counts, which focuses on identifying transcripts from which the reads could have originated based on a human k-mer reference library generated by Kallisto. The platform-generated gene counts were used for downstream analysis.

### Gene ontology

Gene ontology (GO) analysis was performed utilizing the BioJupies cloud-based application and Jupyter Notebooks for RNAseq data analysis [122]. Gene count tables for each THEK cell line (two control cell lines and two HET cell lines) produced via Elysium were uploaded to the platform. Enrichment analysis was performed utilizing Enrichr [123], which pulls the 500 highest and lowest gene expression signatures between experimental groups (controls vs. HETs). “Biological process” enrichment analysis was performed on the platform and identified statistically over- and under-represented GO terms. Statistical significance was determined by using a P-value <0.05 cut-off after applying Benjamini-Hochberg correction. For each biological process, false discovery rate (FDR), adjusted P-value, and z-score were generated. The P-value and FDR for selected top-ranking GO terms were plotted using Prism 9.

### Viral transduction

A retroviral construct used for live imaging of actin (from Dr. Erika Holzbaur, University of Pennsylvania, Philadelphia, PA, USA) was made by sub-cloning the LifeAct-GFP plasmid insert (Addgene #51010) into the retroviral pLZRS vector.

Human embryonic kidney (HEK) 293T retroviral packaging cells (Phoenix cells [124]; from Dr. Garry Nolan, Stanford University, Palo Alto, CA, USA) or 293FT lentiviral packaging cells (from Dr. Erika Holzbaur, University of Pennsylvania, Philadelphia, PA, USA) were grown in complete DMEM. Transfections used 4 μg retroviral plasmid DNA or 2 μg lentiviral plasmid DNA plus 1 μg psPax2 (Addgene, Cat. #12260) plus 1 μg pMD2.G (VSV-G; Addgene, Cat. #12259). Plasmid DNA was mixed with 12 μL FuGENE 6 (Promega, Cat. #E2691) in 800 μL Opti-MEM (Thermo-Fisher Cat. #31985070) at room temperature for 10 min. The cocktail was added dropwise onto 60-mm dishes of sub-confluent Phoenix or 293FT cells, followed by incubation at 37°C.

After 24 h, collected viral supernatants were centrifuged at 200 g for 5 min to pellet dislodged cells. Aliquoted supernatants were snap-frozen in liquid nitrogen then stored at -80°C. For transduction, keratinocyte native medium was replaced with viral supernatant plus 4 μg/mL polybrene (Sigma, Cat. #H9268) at 37°C for 1 h. After removing the viral medium and a PBS rinse, native medium was replaced and transduced cells were propagated.

### Mechanical dissociation assay

Dispase-based mechanical dissociation assays were performed as previously described [125]. Keratinocytes were plated at a density of 1.5 x 10^6^ cells per well of 6-well cell culture dishes. Once confluent, the calcium concentration of the medium was adjusted to 1.3 mM. Vehicle control (DMSO) or chemical inhibitors were added as described in each figure legend. After 18-24 h, monolayers were washed with PBS and then incubated with 500 μL dispase (5 U/ml) in Hank’s balanced salt solution (Stemcell Technologies, Cat. #07913) for 25 min at 37°C. Then, 4.5 ml PBS was added to the wells and released monolayers plus 5mL of diluted dispase were transferred into 15 mL conical tubes. The tubes were placed together in a rack and inverted 2-10 times to induce mechanical stress. Monolayer fragments were transferred back into 6-well cell culture plates and imaged with an iPhone 14, 48-megapixel digital camera. Fragment counts were generated by hand or by examining images in FIJI.

### Organotypic epidermal cultures

Organotypic human epidermal “raft cultures” were generated as previously published [26, 40]. Cultures grown from NHEKs were differentiated in E-medium, a 3:1 mixture of DMEM:Ham’s F12 (Thermo-Fisher, Cat. #11765054) supplemented with 10% FBS, 180 µM adenine (Sigma Cat. #A2786), 0.4 µg/mL hydrocortisone (Sigma, Cat. #H0888), 5 µg/mL human insulin (Sigma Cat. #91077C), 0.1 nM cholera toxin (Sigma, Cat. #C8052), 5 µg/mL apo-transferrin (Sigma Cat. #T1147), and 1.36 ng/mL 3,3′,5-tri-iodo-L-thyronine (Sigma Cat. #T6397). Cultures grown from THEKs were differentiated in CellnTEC Prime Epithelial 3D Medium (Zen-Bio, Cat. #CnT-PR-3D).

Dermal collagen rafts were made from J2-3T3 murine fibroblasts in transwells (Corning Cat. #353091). For each raft, 1 x 10^6^ fibroblasts were counted (Thermo Countess3, Cat. #AMQAX2000) and resuspended to 10% of the desired final volume using sterile reconstitution buffer (1.1 g of NaHCO_3_ and 2.39 g of HEPES in 50 mL 0.05 N NaOH). An additional 10% of the final desired volume of 10x DMEM (Sigma-Aldrich Cat. #D2429) was added. High-concentration rat tail collagen I (Corning, Cat. #CB354249) was then added (final concentration 4 mg/mL) and sterile water was used to dilute to the final volume, 2 mL total per raft. The collagen/fibroblast slurry was inverted to mix, then aliquoted as 2 mL per transwell insert placed within a deep 6-well culture plate (Corning, Cat. #08-774-183). The rafts were polymerized for 1 h at 37°C, after which they were submerged in complete DMEM to incubate overnight in the 37°C incubator.

After 24 h, DMEM was aspirated from both upper and lower transwell chambers and 1-2 x 10^6^ keratinocytes were seeded onto each raft in 2 mL of DMEM; additional DMEM was added to the bottom chamber to keep the raft submerged in liquid. Cultures were then incubated overnight at 37°C. The next day, DMEM was carefully removed from both transwell chambers and keratinocytes were placed at an air-liquid interface by adding either CnT 3D medium (for THEKs) or E-medium (for NHEKs) only into the lower chamber until reaching the bottom of the raft. Vehicle control (DMSO) or chemical inhibitors were diluted in the lower chamber medium as described in the figure legends. Organotypic cultures were matured for 5-12 days, replacing medium in the lower chamber every 2-3 days, followed by live imaging or fixation. For histology, the transwell was transferred into a 6-well culture plate and submerged in 10% neutral-buffered formalin (Fisher Scientific, Cat. #22-026-435) for at least 24 hr.

### Tissue processing and histology

Organotypic epidermis was processed for histologic examination by Core A of the Penn SBDRC or the Experimental Histopathology Core of the Fred Hutchinson Cancer Center. Paraffin-embedded formalin-fixed tissue cross-sections of organotypic epidermis or skin biopsies were processed for histology and stained with hematoxylin and eosin (H&E) using standard methods. H&E-stained glass slides were imaged on an EVOS FL imaging system (Thermo-Fisher) using an EVOS 40X long working distance, achromatic, phase-contrast objective (Thermo-Fisher). Images were captured using the embedded high-sensitivity interline CCD color camera.

### Immunoblotting

Whole-cell lysates of keratinocytes or organotypic cultures were made using urea sample buffer (USB) [8 M Urea, 1% SDS, 10% glycerol, 0.0005% pyronin-Y, 5% β-mercaptoethanol, 60 mM Tris, pH 6.8] and were homogenized using a microtip probe sonicator (Fisher Scientific). Lysates were separated by electrophoresis at 80V for 1 h in NuPAGE MES-SDS Running Buffer (Thermo-Fisher, Cat. #NP0002) and loaded into NuPAGE 12% Bis-Tris Gels (Thermo-Fisher, Cat. #NP0343BOX). Proteins were transferred for 60 min at 50 V onto Immobilon-FL membrane (Millipore Cat., IPFL85R) using transfer buffer (25 mM Tris, 192 mM glycine, 20% (v/v) methanol). Membranes were dried overnight. The next day, blots were re-wetted in 100% methanol and re-hydrated in Tris-buffered saline (TBS). Revert total protein stain and Revert wash solution (LI-COR, Cat. #926-10016) were used for total protein staining captured on an Odyssey M Imaging System, followed by incubation in Revert destaining solution and rinsing in water. Membranes were blocked for 60 min at room temperature in Intercept PBS blocking buffer (LI-COR, Cat. #927-70003), then were probed at 4°C overnight in primary antibodies in Intercept PBS blocking buffer with gentle rocking. Blots were washed three times in 1x TBS with 0.1% (v/v) Tween-20 (TBS-T), then incubated 1 hr at room temperature with gentle rocking in dark boxes in Intercept TBS blocking buffer with 0.02% Tween-20 and 0.01% SDS plus fluorescent secondary antibodies (LI-COR) at 1:10,000. Blots were washed three times in TBS-T then scanned on the Odyssey M.

### Immunoblot quantification

Analysis of immunoblots was performed in FIJI (Version 2.16.0/1.54p) software. Fluorescent immunoblotting images produced via Odyssey M are imported into FIJI. When multiple secondary antibodies were used, the channels were split into individual images and all images were converted to 8-bit, black and white images. LUTs were inverted to ensure the protein bands were displayed as dark signal on a light background.

The rectangle tool was selected from the toolbar and the protein band was traced in the first lane for analysis, ensuring the entire protein band was selected. The first lane was selected utilizing *Analyze > Gels > Select first la*ne, the rectangle box was moved to the second lane and selected via *Analyze > Gels > Select next lane*. This process was repeated for all lanes. Once all lanes were designated, *Analyze > Gels > Plot lanes*, produced an intensity profile plot for each lane. Next, we utilized the straight-line tool to draw a straight line across the base of each lane’s curve, generating a “closed peak” to remove background. The wand tool was used to measure the area under each curve by clicking inside of each lane’s closed curve. Quantified values were copied into an accompanying Excel file and each protein analyzed was normalized to a loading control such as GAPDH.

### Fluorescent immunostaining of cells

Keratinocytes were grown on 35 mm glass-bottom cell culture dishes to confluency (MatTek #P35G-1.5-20-C). For staining plakoglobin and desmosomal proteins, cells were fixed in ice-cold 100% methanol at -20°C for 2 min, allowed to dry, then re-hydrated in PBS. For staining other proteins, cells were fixed in 4% paraformaldehyde for 10 min at 37°C. Fixed cells were incubated 30 min at 37°C in blocking solution [0.5% (w/v) bovine serum albumin (BSA, Sigma, Cat. #A9647), 10% (w/v) normal goat serum (NGS, Sigma, Cat. # G9023) in PBS]. The 35-mm plates were then washed with PBS. Primary antibodies were diluted in 0.5% (w/v) BSA in PBS and incubated on the cells overnight at 4°C. Primary antibody dilutions were as follows: mouse anti-Desmoglein 1 (1:100), mouse anti-Desmoglein 2 (1:100), mouse anti-Desmoglein 3 (1:100), mouse anti-plakoglobin/γ-Catenin (1:400), mouse phospho-myosin light chain 2 (1:200). Cells were washed three times with PBS, then incubated with species-specific secondary antibodies diluted at 1:300 (with or without Hoechst at 1:500) in 0.5% (w/v) BSA in PBS for 30 min at 37°C. Cells were washed three times with PBS, then kept in PBS for imaging using spinning-disk confocal microscopy, as detailed below.

### Fluorescent immunostaining of tissues

Formalin-fixed paraffin-embedded tissue cross-sections on glass slides were baked at 65°C for a minimum of 2 hr. Sections were prepared for staining by immersion in 3 baths of xylenes (Fisher) for 3 min each, followed by 3 baths of 95% ethanol for 5 min each, then 70% ethanol for 3 minutes, and finally 3 baths of PBS for 3 min each. Slides were then submerged in antigen retrieval buffer [0.1 M sodium citrate (pH 6.0) with 0.05% (v/v) Tween-20] and heated to 95°C for 15 min. Slides were allowed to cool to room temperature, then rinsed once in PBS. A hydrophobic barrier pen was used to encircle tissue sections. Tissue sections were incubated in blocking buffer [0.5% (w/v) BSA, 10% (v/v) NGS in PBS] within a humidified chamber for 30 min at 37°C. Slides were rinsed for 3 min in each of 3 PBS baths, then incubated overnight at 4°C in primary antibodies diluted in 0.5% (w/v) BSA in PBS in a humidified chamber. Primary antibody dilutions were as follows: mouse anti-desmoglein 1 (1:50), rabbit anti-plakoglobin/ γ-Catenin (1:400), mouse anti-β-Actin (C4, 1:100), and mouse phospho-myosin light chain 2 (1:400). Slides were then washed in 3 baths of PBS for 3 min each then were incubated for 60 min at 37°C in secondary antibodies (+/- Hoechst) in 0.5% (w/v) BSA in PBS in a humidified chamber. Slides were washed in 3 baths of PBS for 3 min each. Finally, Prolong Gold (Thermo-Fisher, Cat. #P36934) was applied to cover the tissue sections under a #1.5 glass coverslip. After drying, slides were held in the dark prior to imaging on a confocal microscope.

### Confocal fluorescence microscopy

A Yokogawa W1 spinning-disk confocal (SDC) system on a Nikon Ti2 microscope with a Hamamatsu ORCA-FusionBT sCMOS camera was used for image acquisition. Samples were illuminated using laser excitation (405, 488, 561, 640 nm) and emitted fluorescence was captured through a 40x 0.95 NA air objective or 60x 1.2 NA water objective (Nikon) and standard filters. For confocal imaging of live submerged cultures, cells transduced with fluorophore-tagged constructs were seeded into 35-mm glass-bottom dishes at least 24 h prior to imaging in their native medium at 37°C in a stage-top environmental chamber with a sliding lid (Okolab).

### ERK biosensor imaging and analysis

For live imaging of the ERK biosensor, THEKs were transduced with ERK-KTR-mClover. HEK293FT cells were transfected with 4 µg pLenti-CMV-PuroDEST-ERK-KRT-mClover DNA (Addgene #59150) plus 12 µl FuGENE 9 (Promega # E2691) in 800µL of Opti-MEM (Thermo Fisher Scientific #31985070) and cultured overnight in DMEM. Lentiviral supernatants were collected from the HEK292FT the next morning and polybrene (Sigma-Aldrich #H9268) was added (4µg/mL). KSFM was removed from THEKs and replaced with viral medium for 1 h at 37°C. Cells were then washed in PBS and placed back in KSFM and expanded.

ERK-KTR-transduced cells were seeded into 35 mm glass-bottom dishes and grown to confluency, then were placed in M154 with 0.07 mM calcium the night before imaging. The next morning, cells were spiked with high calcium (1.3mM) and imaged after 4 hr as described below. Analysis was performed using FIJI, as previously described [41]. The nuclear and cytoplasmic regions for each cell were selected utilizing the polygon selection tool and the “Measure” function was used to calculate mean fluorescence intensity of each region. The nuclear intensity was subtracted from the cytoplasmic intensity and the ERK activity index was calculated as the ratio of cytoplasmic-to-nuclear intensity after background correction.

### Fluorescent immunostaining quantification

Immunostained cell sheets or tissue sections were imaged using confocal microscopy as above. Fluorescence microscopy images were analyzed using FIJI (Version 2.16.0/1.54p). Quantification of fluorescence intensity was performed in a blinded manner using non-visibly identifiable microscopy images. The FIJI “Measure” function was used to calculate the mean intensity across the entire field of fixed cells or a region of interest (ROI) encompassing the entire epithelium, circumscribed using the polygon selection tool. The mean background fluorescence intensity was subtracted from each value. Fluorescence intensity for each protein of interest was normalized to a control protein or Hoechst nuclear stain. The mean fluorescence intensity from each image was averaged for each condition and in multiple experimental replicates for statistical analysis; the average intensity of pooled control samples was normalized to 1.

Line scan analysis was completed in FIJI utilizing the “Straight Line” function for multiple images per experiment; the same line position was used for all images. The “ROI Manager” was used to add the line as a measurable region of interest and the “Measure Plot” tool was used to document the fluorescence intensity along each pixel. Graphs were generated using PRISM; the x-axis plots each pixel location and the y-axis plots the fluorescence intensity; representative line-scans were depicted.

### Junctional actin quantification

To quantify actin recruitment to cell-cell junctions, polymerized actin was measured using phalloidin staining to generate intensity profiles in FIJI (Version 2.16.0/1.54p). Lines were drawn perpendicular to junctions from cytoplasm to neighboring cytoplasm; 15 pixel-wide lines were used to avoid sampling artifacts and increase the signal-to-noise ratio. Each line generated a profile of intensity over distance as measured in microns. Two measurements were made per junction when possible and junctions were chosen based on monolayer integrity, avoiding junctions from cells compromised during fixation or junctions on field-of-view edges. A Gaussian was fitted to each intensity profile to extract the full width at half maximum (FWHM).

### Statistics

Data graphing and statistical analyses were performed in Prism version 9 (GraphPad). Sample size, center definition, dispersion measures, and statistical tests are included in figure legends. Means of 2 normally distributed groups were compared using a 2-tailed unpaired or paired Student’s t-test. Means from ≥2 normally distributed groups were compared by 1-way ANOVA with P-values adjustment for multiple comparisons. P-values <0.05 were deemed statistically significant. Graphs include exact *P* values.

### Study approval

Normal human epidermal keratinocytes (NHEKs) from neonatal foreskins and human skin biopsy sections were provided by the Penn Skin Biology and Diseases Resource-based Center (SBDRC) in a de-identified fashion under protocols (#808224; #808225) approved by the University of Pennsylvania Institutional Review Board (IRB). Use of de-identified cells and tissues (collected for clinical purposes that would otherwise be discarded) was deemed exempt by the IRB for written informed consent.

## AUTHOR CONTRIBUTIONS

Conceptualization: J.L.A., C.L.S. Data Curation: J.L.A., A.P., A.T., C.L.S. Formal analysis: J.L.A., K.P., A.A., M.C.M., K.S., C.L.S. Investigation: J.L.A, A.P., A.T., C.J.T., M.C.M., N.S., M.K.S., C.L.S. Resources: J.E.G., C.L.S. Supervision: C.L.S. Visualization: J.L.A, A.P., A.T., N.S., C.L.S. Validation: J.L.A., M.K.S., K.S., C.L.S. Writing (original draft): J.L.A., C.L.S. Writing (review and editing): J.L.A, C.L.S.

## ACKNOWLEDGEMENTS

We thank the NIH-funded Skin Biology and Diseases Resource-Based Centers at Univ. of Pennsylvania (P30AR069589) and Univ. of Michigan (P30AR075043) for keratinocytes, CRISPR/Cas9 editing, histology, and biopsy sections. We thank Dr. Mary Regier of the UW ISCRM Genomics Core for assistance with RNA sequencing. C.L.S. was supported by NIH (K08AR075846, R03AR082896, R03TR005428) and the LEO Foundation (LF-OC-23-001393, LF-SE-25-800075). J.L.S. was supported by the Univ. of Washington (UW) Institute of Translational Health Sciences funded by NIH (TL1TR002318). A.P. and. A.T. were supported by the Foundation for Ichthyosis and Related Skin Types and an Innovation Pilot Award from the UW Institute for Stem Cell and Regenerative Medicine. M.C.M. was supported by the UW School of Medicine-Gonzaga University Health Partnership. C.J.T. and N.S. were supported by the Dermatology Foundation. J.E.G. and M.K.S. were supported by NIH (P30AR075043) and the Taubman Medical Research Institute. K.S. was supported by the M.J. Murdock Charitable Trust. BioRender.com was used for the graphical abstract and figure illustrations.

